# GPU handling of OPC water in GROMACS: validation by protein folding and a 1.6-fold speed-up

**DOI:** 10.64898/2026.11.19.767864

**Authors:** Takashi Kato, Nobuhiro Shojima, Toshimasa Yamauchi

## Abstract

Molecular dynamics (MD) simulation is one of the core computational methods of the life sciences, used to study protein folding, receptor activation and drug binding in atomic detail. The timescales it can reach depend on speed, and running each step entirely on a graphics processing unit (GPU) is now essential for fast simulation. Optimal Point Charge (OPC) water, recommended for the ff19SB protein force field, carries its negative charge on a virtual site, and in GROMACS 2026.3 virtual sites prevent the coordinate update from running on the GPU. The GROMACS developers have recently added GPU handling of virtual sites to the development version as an experimental feature. Whether simulations run in this way agree with the established CPU calculation has to be measured, and no such test has been reported over microseconds or in a membrane-protein system of production size. We compared a modified build that handles OPC sites within the GPU update (Optimized) with the official release (Reference). The two builds placed the charge site with the same error (2.3 *×* 10^*−*7^ nm) and gave relative force differences below 10^*−*6^. In five matched pairs of 5 *µ*s simulations, the Trp-cage miniprotein folded with both builds (three of five Reference and five of five Optimized runs; *P* = 0.44), and the folded ensembles matched. Both builds reproduced the density, diffusion coefficient, dielectric constant and second hydration shell that distinguish OPC from TIP3P water. In a 166,826-atom membrane system of the glucagon-like peptide-1 receptor, residue hydration agreed between the builds (*r* = 0.99), and Optimized ran 1.57–1.66 times as fast as tuned Reference on two GPUs, with 87–97% of the time saved coming from moving the coordinate update to the GPU. In these tests, GPU handling of OPC sites gave a substantial speed gain with no sign of altered physical behaviour.

## 1 Introduction

Molecular dynamics (MD) simulation has become one of the core methods of the life sciences [1, 2, 3]. By following the motion of every atom in a protein and its surroundings, it has shown how proteins fold [4, 5], how G-protein-coupled receptors change conformation on activation [6] and how drugs bind their targets [7], and it is now used routinely alongside structural biology and drug discovery. Many of these processes take microseconds or longer, whereas a simulation advances in steps of a few femtoseconds [2]. What MD can show therefore depends directly on how many steps can be computed per day. Since MD programs were moved to graphics processing units (GPUs) [8, 9, 10, 11], that number has been determined largely by how much of each step runs on the GPU.

Water accounts for most of the atoms in a typical simulation, and the choice of water model influences the result [12]. Three-site models such as TIP3P place the negative charge on the oxygen [13]. Four-site models, including TIP4P-Ew [14], TIP4P/2005 [15] and Optimal Point Charge (OPC) water [16], move it to an additional massless site and reproduce the properties of liquid water more closely. OPC has been reported to improve simulations of intrinsically disordered proteins [17], and the authors of the ff19SB protein force field recommend it after comparing explicit water models [18]. In OPC the additional site, M, is a virtual site on the H–O–H bisector: at every step the program must construct its position from the three atoms and, once the forces have been computed, redistribute the force acting on M back to those atoms.

GROMACS is one of the most widely used MD programs [19]. It can run an entire step on the GPU, including the coordinate update [11], but release 2026.3 cannot do so when the system contains virtual sites [20, 21]. With OPC water, the coordinate update, the constraints and the site operations therefore run on the central processing unit (CPU). Coordinates and forces then have to be exchanged between CPU and GPU at every step, and this synchronization slows the simulation. We meet this limitation in our own work on the glucagon-like peptide-1 receptor (GLP-1R), a class B G-protein-coupled receptor and a drug target in type 2 diabetes and obesity [22, 23]: our simulations of GLP-1R and its R131Q variant, 38 membrane runs totalling 22.8 *µ*s, used OPC water [24]. The more accurate water model is thus the slower one to run.

The GROMACS developers are working on this problem. They have demonstrated a prototype that constructs virtual sites and spreads their forces on the GPU, using a box of 395 four-site TIP4P waters [25], and in November 2026 they merged these operations into the GPU update of the development version, as an experimental feature [26]. Moving the update to the GPU leaves the equations, the precision and the time step as they are and changes only where they are evaluated. As with any change of hardware, the numbers are still not reproduced digit for digit: floating-point sums depend on the order of their terms [27], GPU code sums forces in a different order from CPU code [28], and because MD is chaotic, two trajectories started from the same state separate within picoseconds [29]. Whether the modified calculation is correct therefore has to be established from the results that a simulation is run for. For GPU-handled virtual sites, this has been reported neither for long simulations nor for a biological outcome.

This study provides that validation. We compare a modified build of GROMACS 2026.3 that handles OPC sites within the GPU update (Optimized) with the official release (Reference), and ask two questions: whether the modified build gives the same physical and biological results, and how much faster it is. For the first question we examine the position of the site and the forces in a single step, and then two results that emerge only over long simulations: the folding of a protein and the properties of OPC water. As the folding test we chose Trp-cage, a 20-residue miniprotein that is one of the most widely studied models of protein folding [30, 5]. Folding suits the purpose: it emerges only after hundreds of millions of steps, and its end point can be checked against experiment. For Trp-cage the native structure is known [31], folding takes about 4 *µ*s in experiments [32] and is thus within reach of simulation, the stability of the fold depends on the water model [33], and folding with ff19SB has been reported [34]. For the second question we measure the speed of a membrane system from our own research, GLP-1 bound to the wild-type GLP-1R [24].

## 2 Results

### 2.1 OPC charge-site geometry and solvent structure

We first tested whether the GPU code places the OPC site where the model defines it. At the end of each of the ten Trp-cage trajectories, we recomputed the position of M from the oxygen and hydrogen coordinates of every water molecule, using the construction rule of the topology, and measured its distance from the position of M that the simulation itself had written to the same checkpoint. The root-mean-square (RMS) distance over the 9,300 waters was 2.31 *×* 10^*−*7^ nm with Reference and 2.32 *×* 10^*−*7^ nm with Optimized (Figure 1B). The difference between the group means, 7 *×* 10^*−*10^ nm, was not significant (Welch’s *t*-test, *P* = 0.27; five trajectories per build). An error of this size is expected from single-precision storage of the coordinates.

**Figure 1:**
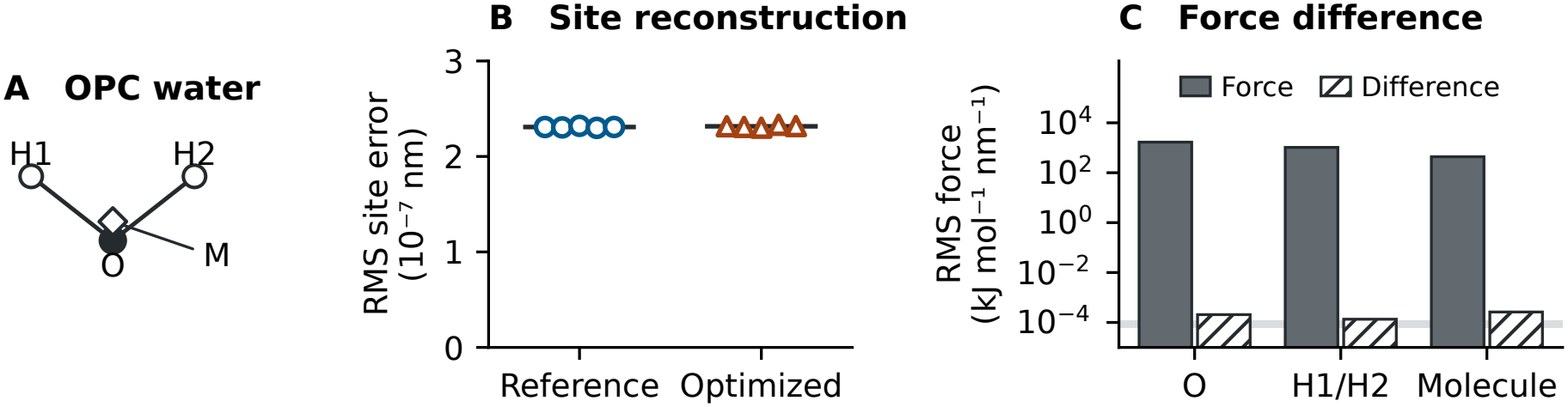
OPC site geometry and forces. (A) Charge site M. (B) RMS distance between the M position recomputed from the O and H coordinates and the M position stored by the simulation, for 9,300 waters per endpoint. Circles show Reference, triangles Optimized and bars the means. (C) RMS force in Reference (filled) and RMS difference between the two builds (hatched) on a common input, for oxygen, hydrogen and the net force per water molecule, on a logarithmic scale. The grey band marks the spacing of single-precision numbers at these force magnitudes.

We then compared the forces that the two builds compute for one common starting configuration (Figure 1C). The RMS forces on oxygen, on hydrogen and on whole water molecules were about 1,680, 1,030 and 440 kJ mol^*−*1^ nm^*−*1^. The two builds differed by an RMS of 2.1 *×* 10^*−*4^, 1.3 *×* 10^*−*4^ and 2.6 *×* 10^*−*4^ kJ mol^*−*1^ nm^*−*1^, about seven orders of magnitude less. A difference of this size is the last digit that single precision can represent for forces of this magnitude, as expected when the same sum is accumulated in a different order. In relative terms the differences were 1.23 *×* 10^*−*7^, 1.30 *×* 10^*−*7^ and 6.02 *×* 10^*−*7^; the value is larger for whole molecules only because the forces on the sites of one molecule largely cancel, so that a similar absolute difference is divided by a smaller total. Every M site had zero residual force after redistribution.

If the site or its forces were handled incorrectly, the structure of the water itself would change. We therefore compared intermolecular water–water pair distributions at the end of each trajectory. The oxygen–oxygen and hydrogen–M distributions of the two builds coincided, and the first hydrogen–M peak, which reports the hydrogen bond to the charge site, lay at 0.205–0.210 nm in both (Figure S1, Table S1).

### 2.2 Trp-cage reached a native-like fold in both implementations

We next asked whether the build reproduces a biological process that depends on the solvent. We started one Reference and one Optimized trajectory from each of five independently equilibrated, non-native structures and followed them for 5 *µ*s. A frame was counted as native-like when it met four criteria at once: backbone fit to the nuclear magnetic resonance (NMR) structure, native contacts, packing of tryptophan 6 against prolines 12 and 18, and formation of the N-terminal helix (Methods). All five Optimized trajectories reached a persistent native-like fold, as did three of the five Reference trajectories (Figure 2; C*α* deviations from the NMR structure are shown in Figure S2). This difference in the number of folded trajectories is not significant (Fisher’s exact test, *P* = 0.44).

**Figure 2:**
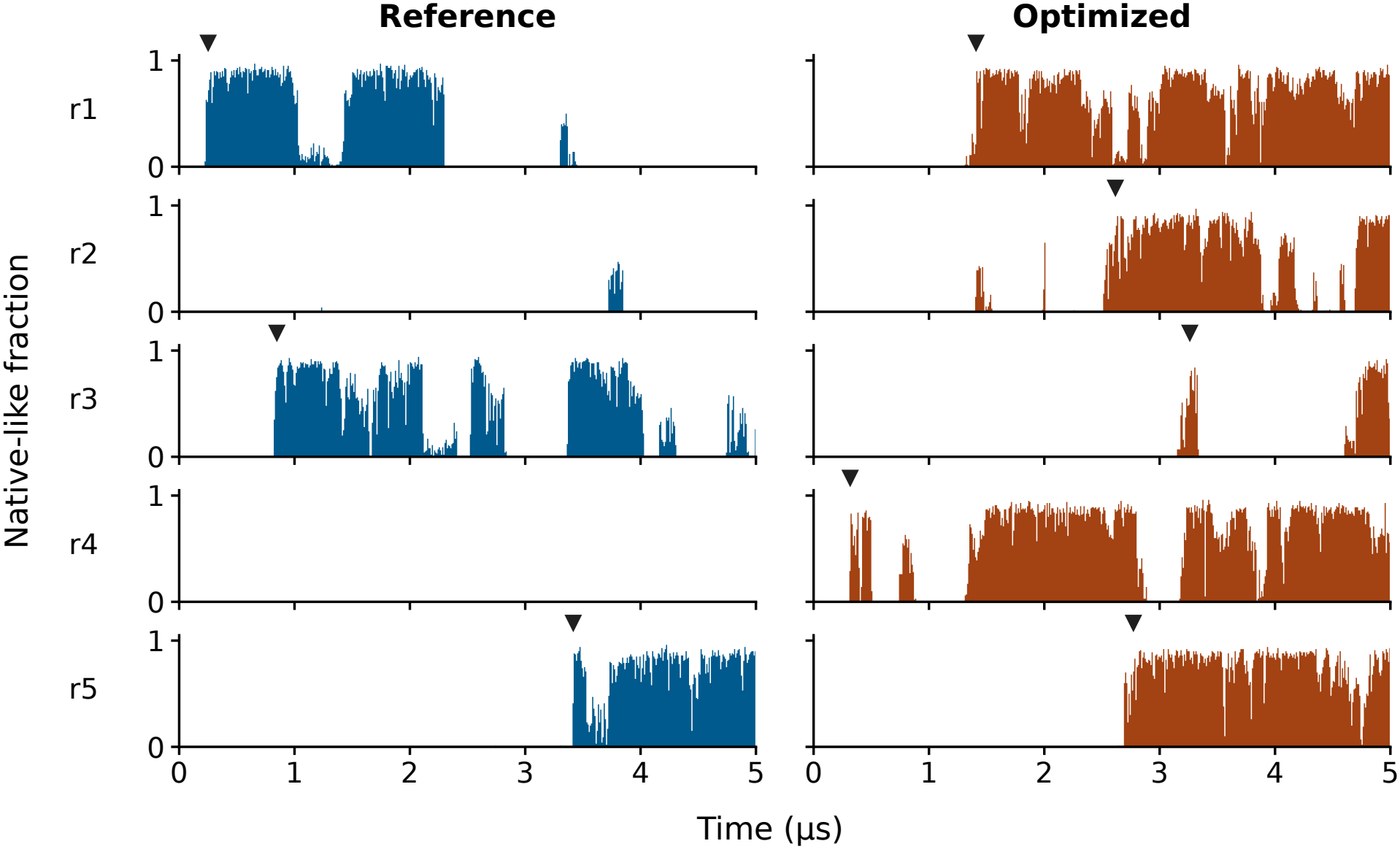
Folding of Trp-cage. Each panel shows the fraction of frames, in consecutive 10-ns windows, that met all four native-like criteria over 5 *µ*s. Each row shows the two runs started from the same input. Triangles mark the first confirmed persistent native-like event.

The times of first entry into the fold overlapped between the builds (0.25–3.42 *µ*s with Reference and 0.32–3.26 *µ*s with Optimized; Table S2; log-rank test, *P* = 0.18) and are of the order of the experimental folding time of about 4 *µ*s [32]. If folding is a single-exponential process with a time constant near that value, a 5 *µ*s trajectory has roughly a 30% chance of not folding at all, so two Reference trajectories without a persistent fold are not unexpected.

Once folded, the peptide adopted the same structure with both builds. In the last native-like frame of each folded trajectory, the C*α* deviation from the NMR structure was 0.08–0.21 nm with Reference and 0.07–0.18 nm with Optimized, with tryptophan 6 packed against prolines 12 and 18 (Figure 3A). Over all native-like frames, about 43,000 with Reference and 88,000 with Optimized, the distributions of backbone deviation, native contacts and the two tryptophan–proline distances coincided (Figure 3B), with 93–99% overlap between the builds. The trajectory means agreed as well (Table 1): the mean C*α* deviation was 0.10 nm with both builds, and the fraction of native contacts 0.87. The only nominally significant difference among five quantities, in the tryptophan 6–proline 18 distance, amounts to 0.0007 nm.

**Table 1:** The folded state in the two builds. Values are the mean *±* sample standard deviation of trajectory means over native-like frames (three Reference and five Optimized trajectories). *P* values are from Welch’s *t*-test and are not corrected for multiple comparisons.

| Quantity | Reference | Optimized | *P* |
|---|---|---|---|
| Cα RMSD from NMR structure (nm) | 0.104 ± 0.010 | 0.101 ± 0.005 | 0.61 |
| Fraction of native contacts | 0.874 ± 0.009 | 0.868 ± 0.008 | 0.40 |
| Trp6–Pro12 distance (nm) | 0.3727 ± 0.0006 | 0.3726 ± 0.0007 | 0.84 |
| Trp6–Pro18 distance (nm) | 0.3681 ± 0.0001 | 0.3688 ± 0.0005 | 0.03 |
| Helical residues among residues 2–8 | 6.76 ± 0.04 | 6.76 ± 0.05 | 0.80 |

**Figure 3:**
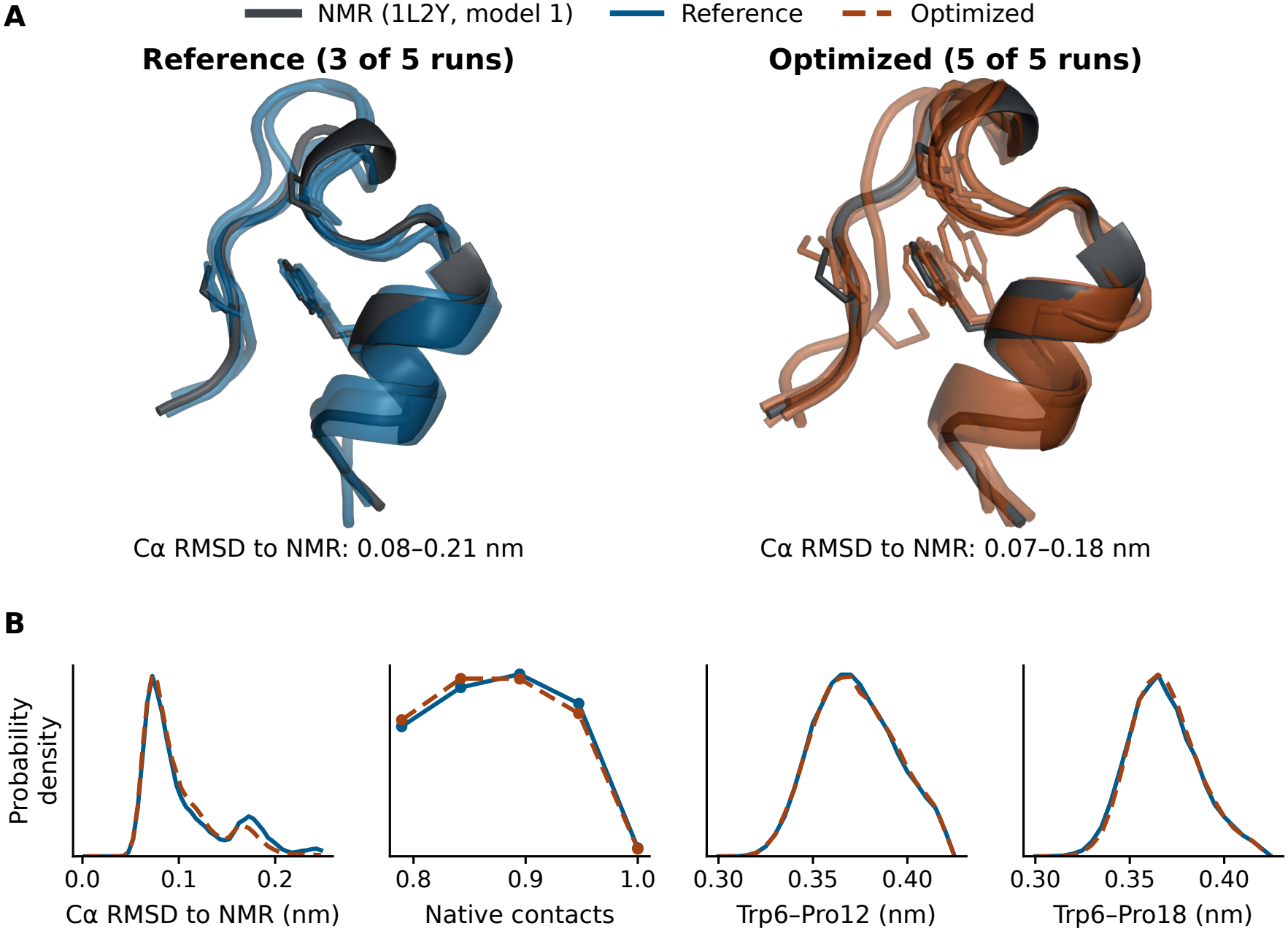
The folded state of Trp-cage in the two builds. (A) The last native-like frame of every trajectory that reached a persistent fold (colour) is superposed on NMR model 1 of 1L2Y (grey) by its C*α* atoms. Sticks show tryptophan 6, proline 12 and proline 18. (B) Distributions over all native-like frames of those trajectories: C*α* RMSD from the NMR structure, fraction of native contacts, and the minimum heavy-atom distances from tryptophan 6 to proline 12 and proline 18. The ranges are bounded by the criteria that define a native-like frame.

Optimized did not fold less than Reference by any measure. The fraction of the 5 *µ*s spent in native-like structures varied widely among the runs of each build, from 0 to 35% with Reference and from 6 to 54% with Optimized (Table S2), because it depends mainly on how early a run happened to fold. The two builds gave similar values (Welch’s *t*-test, *P* = 0.16).

### 2.3 Water behaviour that distinguishes OPC was preserved

The tests above compare the two builds with each other. The reason for using OPC is the behaviour of its water, so we next tested whether Optimized reproduces that behaviour. We simulated a box of 9,300 water molecules for 10 ns with three set-ups: TIP3P, the common three-point model [13], with Reference; OPC with Reference; and OPC with Optimized. The properties in which OPC differs from TIP3P were reproduced by both builds (Table 2). TIP3P water diffused 2.5 times as fast as OPC water, was 11 kg m^*−*3^ less dense, had a dielectric constant near 100 and showed almost no second hydration shell in the oxygen–oxygen radial distribution *g*_OO_ (Figure 4A). With OPC, Reference and

**Table 2:** Bulk properties of water at 298 K and 1 bar. Values are the mean *±* sample standard deviation of five consecutive 2-ns blocks of one 10-ns simulation of 9,300 molecules. *P* values compare OPC Reference with OPC Optimized (Welch’s *t*-test, uncorrected). Diffusion coefficients are not corrected for the finite size of the box, which was the same in all three simulations. Experimental values are those compiled by Izadi et al. [16].

| Property | TIP3P | OPC Reference | OPC Optimized | *P* | Experiment |
|---|---|---|---|---|---|
| Density (kg m<sup>−3</sup>) | 986.0 ± 0.1 | 997.1 ± 0.1 | 997.1 ± 0.1 | 0.19 | 997.0 |
| Diffusion coefficient (10<sup>−5</sup> cm<sup>2</sup> s<sup>−1</sup>) | 5.76 ± 0.03 | 2.28 ± 0.03 | 2.26 ± 0.01 | 0.14 | 2.30 |
| Dielectric constant | 99 ± 3 | 75 ± 4 | 76 ± 3 | 0.59 | 78.4 |
| *g*<sub>OO</sub>, first peak | 2.70 ± 0.01 | 3.22 ± 0.01 | 3.22 ± 0.01 | 0.89 | – |
| *g*<sub>OO</sub>, second peak | 1.009 ± 0.003 | 1.105 ± 0.001 | 1.105 ± 0.001 | 0.89 | – |

**Figure 4:**
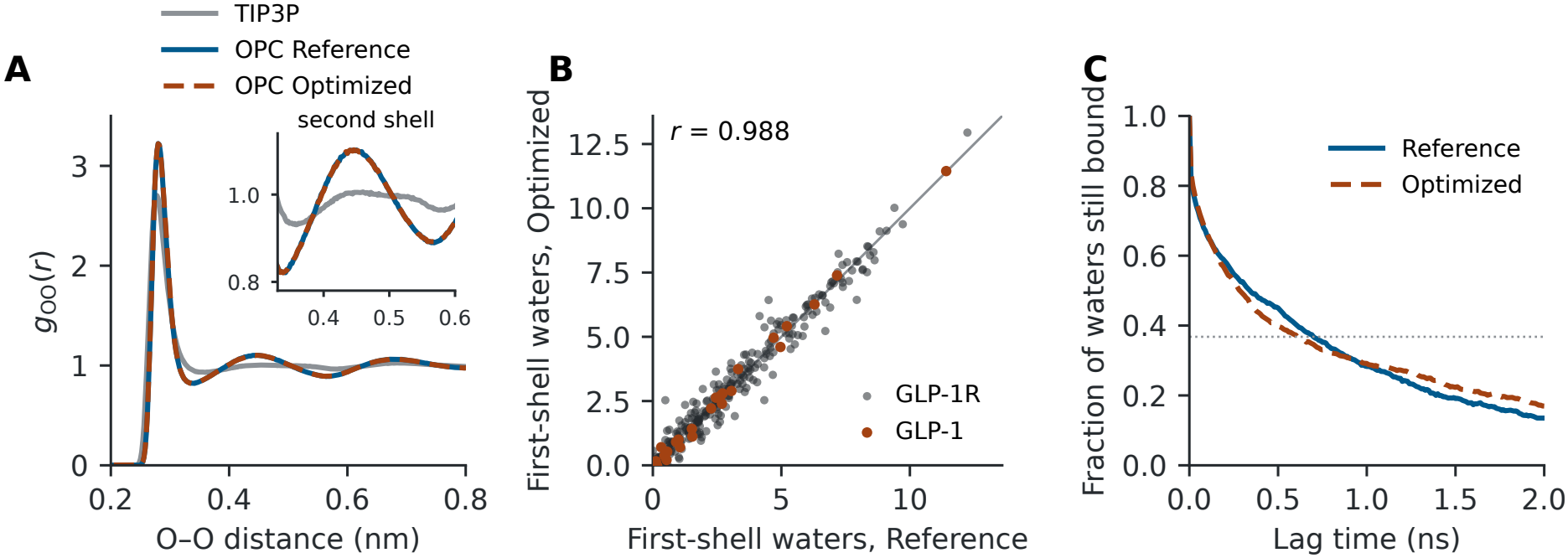
Water behaviour in bulk and at GLP-1R. (A) Oxygen–oxygen radial distribution of pure water for TIP3P, OPC with Reference and OPC with Optimized; the inset enlarges the second hydration shell. (B) Mean number of water molecules in the first hydration shell (oxygen within 0.35 nm of a nitrogen or oxygen atom) of each of the 425 residues of GLP-1 and GLP-1R in 8-ns simulations with the two builds. The line marks equality. (C) Fraction of the water molecules in the first shell of GLP-1 residues His7–Asp15 that are in that shell again after the given lag time. The dotted line marks 1*/e*.

Optimized both gave a density of 997 kg m^*−*3^, diffusion coefficients of 2.28 and 2.26 *×*10^*−*5^ cm^2^ s^*−*1^, close to the experimental 2.30, a dielectric constant near the experimental 78, and the same second shell. None of the five properties differed significantly between the two builds, and for each of them the difference between the builds was small compared with the difference between the water models.

We then asked the same question for water at a protein. Structural water in the peptide-binding pocket contributes to ligand recognition in class B receptors such as GLP-1R [23], so the hydration of the bound peptide is the place where an error in water handling would matter for our own work. Starting from the state that began the published simulation of GLP-1-bound GLP-1R [24], we ran the receptor system for 8 ns with each build and saved all water positions every 10 ps. For each of the 425 residues of GLP-1 and the receptor we counted the water molecules in its first hydration shell. The mean counts of the two builds were highly correlated (*r* = 0.988; Figure 4B). Their RMS difference was 0.39 molecules, no larger than the difference between the first and second halves of a single run (0.43 molecules for Reference and 0.40 for Optimized). The protein as a whole held 897 *±* 16 first-shell waters with Reference and 897 *±* 20 with Optimized (*P* = 0.98, four 2-ns blocks per build), and the N-terminal segment of GLP-1 (His7–Asp15), which inserts into the transmembrane pocket, held 8.6 *±* 0.6 and 8.3 *±* 0.7 (*P* = 0.61). In both builds the receptor residues most often joined to this segment through a shared water molecule were Asn300, Arg380 and Thr298 (Arg380: 84% of frames with Reference and 86% with Optimized). Water molecules bound to the segment were exchanged at similar rates: the fraction still bound fell to 1*/e* after 0.7 ns with Reference and 0.6 ns with Optimized (Figure 4C). The number of waters bridging the segment and the receptor was 5.4 *±* 1.3 and 4.0 *±* 0.3 (*P* = 0.11).

### 2.4 GPU coordinate updates gave the largest speed gain in GLP-1R

The sections above asked whether Optimized computes the same physics as Reference. This section asks a different, practical question: how much faster it is on a system of real size, and where the gain comes from. With all features enabled, Optimized ran the GLP-1R input at 157 and 138 ns/day on the RTX 3080 and RTX 2080 Ti, against 95 and 88 ns/day for tuned Reference, a gain of 1.66- and 1.57-fold (Table 3).

**Table 3:** Throughput of the GLP-1R membrane system. Tuned Reference is compared with Optimized with all features enabled. Values are the mean *±* sample standard deviation of three timed runs. The ratio uses the mean step times.

| GPU | Reference (ns/day) | Optimized (ns/day) | Ratio |
|---|---|---|---|
| RTX 3080 | 94.6 ± 0.0 | 156.8 ± 1.1 | 1.66 |
| RTX 2080 Ti | 88.0 ± 0.9 | 138.4 ± 1.7 | 1.57 |

The build combines four changes, and only the first is specific to OPC water, so the total alone does not show which of them a user or developer needs. The four changes are the following. (i) *GPU update with OPC sites*: integration, constraints, construction of M and redistribution of its force all run on the GPU, so that coordinates need not return to the CPU between steps. (ii) *GPU CMAP*: the correction map (CMAP) is the ff19SB term that adjusts the energy of each residue as a function of its two backbone dihedral angles [18]; while it is evaluated on the CPU, coordinates and forces must still be exchanged with the CPU at every step. (iii) *Fused force redistribution*: the redistribution of the M-site force is merged into the GPU operation that sums the force contributions (force reduction), which saves one pass over the force array. (iv) *CUDA Graphs*: a feature of the CUDA programming interface that records the GPU operations of a step once and then submits them as a single unit instead of one at a time.

To attribute the gain, we enabled these changes one after another in Optimized, in the order given, and timed each configuration (Figure 5). Moving coordinate updates, constraints and OPC site operations together to the GPU reduced step time by about 40% while CMAP still ran on the CPU. In follow-up traces, retaining updated coordinates on the GPU removed approximately one coordinate upload per regular step.

**Figure 5:**
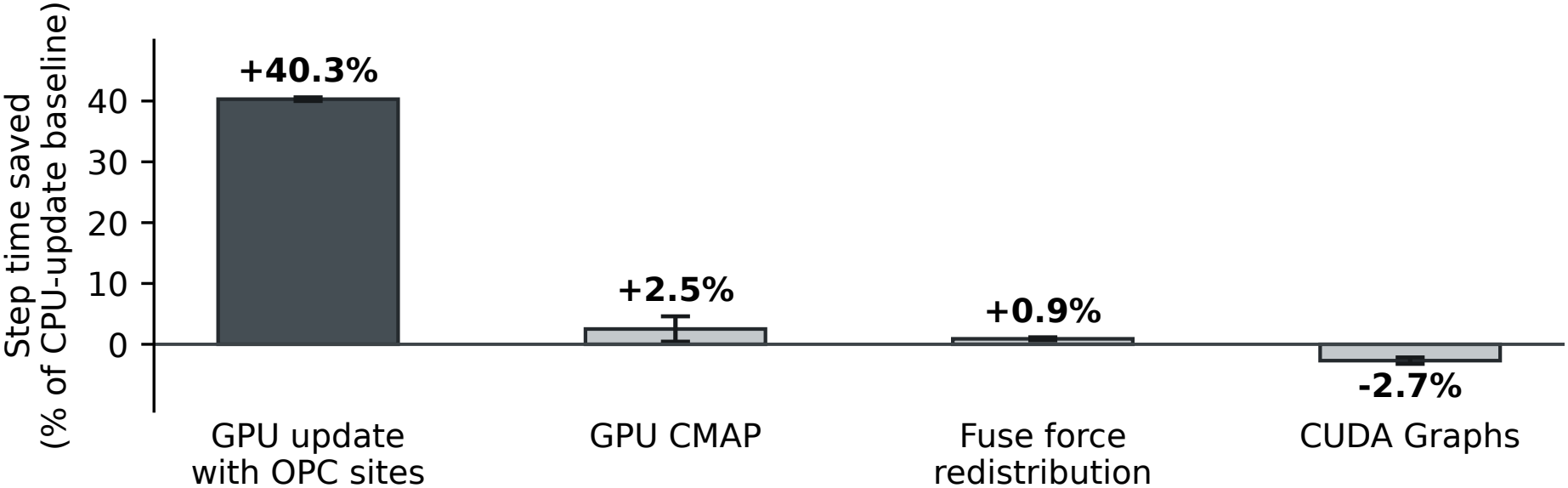
Time saved by the four changes in Optimized, enabled one after another on the GLP-1R system: GPU update with OPC sites, GPU evaluation of the backbone correction map (CMAP), fusion of M-site force redistribution with force reduction, and CUDA Graphs. Each change is measured against the preceding configuration and expressed as a percentage of the same CPU-update baseline on that device. Bars give the mean across the RTX 3080 and RTX 2080 Ti. Capped lines span their two values. Negative savings indicate a longer step.

The other three changes mattered much less. Moving CMAP to the GPU and fusing force redistribution with force reduction gave small gains; on the RTX 2080 Ti, GPU CMAP saved 0–1 s per timed run, which is the resolution of the timer. The coordinate-update stage accounted for 87–97% of the time saved by the first three changes. Enabling CUDA Graphs lengthened the step, so the fastest configuration on both GPUs used the first three changes with CUDA Graphs off.

## 3 Discussion

Handling the OPC virtual site on the GPU made our GLP-1R simulations 1.57–1.66 times as fast, and in the tests reported here we found no sign that it changed the physical behaviour of the system. This is the result we expected, and the reason lies in what the modification does. Optimized computes the same forces from the same equations at the same precision as Reference. It changes the place where the coordinate update is carried out, and thereby removes a round trip between CPU and GPU. A change of this kind can reach the results only through rounding, and each of our observations fits that expectation. The speed gain came almost entirely from the removed transfer: the build is faster because it communicates less, while the amount of computation is unchanged. The forces of the two builds differed in the last digit of single precision, the size expected when one sum is accumulated in a different order, and nothing larger appeared at any level we examined: the position of the site, the properties that distinguish OPC from TIP3P water, the hydration of GLP-1 and its receptor, and the folded state of Trp-cage all agreed. Differences of this size also arise whenever a simulation is moved to different hardware, a different compiler or a different number of threads. We therefore regard GPU handling of the OPC site as a change in how the calculation is executed, comparable to running it on another machine, with the method itself left intact.

Of the four changes in the build, the GPU update with OPC sites accounted for nearly all of the gain: it shortened the step by about 40% on both devices. GPU evaluation of CMAP and fused force redistribution each added less than 3%, and CUDA Graphs lengthened the step, so the fastest configuration ran without them. The part of our build that matters is therefore the one that the GROMACS developers have now merged into the development version [25, 26]. Our results indicate that this approach pays off for OPC water in a receptor system of production size, and they give no indication that it compromises protein folding. We tested our own build only, and the upstream implementation remains to be benchmarked in the same way.

Several limitations apply. Five runs per build are too few to tell whether the builds differ in how often or how early Trp-cage folds. The comparisons of bulk water and of GLP-1R hydration rest on one short simulation per set-up (10 and 8 ns); 8 ns do not fully sample the exchange of the water that bridges GLP-1 and the receptor. The OPC Reference water simulation ran on a different workstation from the other two. The timings come from single-GPU runs with CUDA on two GPU models. Long matched GLP-1R trajectories and a direct comparison with the upstream implementation are the natural next steps.

In summary, a build of GROMACS that keeps OPC water on the GPU through the coordinate update ran a membrane-protein system of production size about 1.6 times as fast as the official release and reproduced its results, from the forces of a single step to the properties of the water and the folding of a miniprotein. For groups that use ff19SB with OPC water in GROMACS, this narrows the gap between the recommended water model and the fastest execution path, and it supports the integration of virtual sites into the GPU update.

## 4 Methods

### Implementation

Optimized is GROMACS 2026.3 with four additions for CUDA GPUs: construction of OPC sites and redistribution of their forces within the GPU update; GPU evaluation of CMAP; fusion of the force redistribution with the GPU force reduction; and optional use of CUDA Graphs. In the benchmark build each of the last three can be switched separately at run time. The Trp-cage production runs used build opcfast20260809g with all four enabled. The source is available at the address given under Data and code availability.

### Simulations

The Trp-cage peptide (TC5b) was simulated in GROMACS [19, 11] with ff19SB and OPC [18, 35]. The system contained 9,300 waters and one chloride. Each matched pair read the same compiled input, including velocities, and ran for 5 *µ*s at 300 K and 1 bar with a 2 fs step. Reference updated coordinates and applied constraints on the CPU. Optimized (`opcfast20260809g`x) ran coordinate updates, constraints, OPC site construction and CMAP on the GPU, with CUDA Graphs enabled. It fused OPC force redistribution with force reduction on compatible steps. Separate force-output runs supplied unrounded initial forces. Site errors of the two builds were compared with Welch’s *t*-test. We computed intermolecular pair distributions from final checkpoints with periodic boundaries and 0.005 nm bins, giving each replica equal weight. The time series stored protein coordinates.

### Water behavior

Pure water (9,300 molecules, cubic box of about 6.5 nm) was built with the TIP3P and OPC parameters distributed with the ff19SB port [35], energy-minimized, equilibrated for 0.5 ns and simulated for 10 ns at 298.15 K and 1 bar (velocity-rescaling thermostat, Parrinello–Rahman barostat, 2 fs step, 1.0 nm cut-offs, particle-mesh Ewald electrostatics), saving coordinates every 1 ps. OPC Reference and OPC Optimized started from the same compiled input. Density, the self-diffusion coefficient (linear fit of the oxygen mean-square displacement between 50 and 500 ps), the static dielectric constant (fluctuation of the total dipole moment) and *g*_OO_ were computed with the GROMACS analysis tools for five consecutive 2-ns blocks. The OPC Reference water simulation ran on a second workstation (AMD Ryzen 9 9950X processor, RTX 5090, GROMACS 2026.3 built with CUDA 12.9); the other two water simulations and all GLP-1R simulations ran on the workstation described under Timing. For GLP-1R we recompiled the published production input [24] from its starting coordinates and velocities so that all atoms were saved every 10 ps, and ran 8 ns with each build on the RTX 3080 (Reference with CPU update, Optimized with GPU update, CMAP on the GPU and CUDA Graphs enabled). A water molecule was assigned to the first hydration shell of a residue when its oxygen lay within 0.35 nm of a nitrogen or oxygen atom of that residue, and was counted as bridging when it belonged to the shells of a residue of GLP-1 His7–Asp15 and of a receptor residue at once. The residence curve is the intermittent autocorrelation of shell membership. Block means (four 2-ns blocks, the common starting frame excluded) were compared with Welch’s *t*-test.

### Fold detection

Frames sampled every 100 ps had to satisfy four criteria: C*α* RMSD *≤* 0.25 nm from NMR model 1 of 1L2Y, native-contact fraction *≥* 0.75, both Trp–Pro distances *≤* 0.42 nm and at least five helical residues among residues 2–8 according to the Kabsch–Sander secondary-structure assignment [36]. A persistent event required at least 80% occupancy in 10 ns and a frame passing the stricter 0.15 nm core-fit threshold in trajectory frames stored every 10 ps. Occupancies use the full 5 *µ*s of each run. For the structural comparison we took the last native-like frame of each trajectory with a persistent event and fitted its C*α* atoms to the NMR model. Folded-state quantities were averaged over the native-like frames of each such trajectory and the trajectory means compared with Welch’s *t*-test. Numbers of folded trajectories were compared with Fisher’s exact test, first-entry times with the log-rank test (trajectories without a persistent event censored at 5 *µ*s) and occupancies with Welch’s *t*-test, treating the five runs of each build as independent. All tests are two-sided.

### Run options

Optimized needs no options beyond those of standard GROMACS. With `gmx mdrun –ntmpi 1 -nb gpu -pme gpu -pmefft gpu -bonded gpu -update gpu -nstlist 150`, the build detects OPC water and handles its sites, CMAP and the fused force redistribution on the GPU, and uses CUDA Graphs. The fastest configuration measured here adds the environment variable `GMX_DISABLE_CUDA_GRAPH=1`, which turns CUDA Graphs off. The options and the variable are the same on Linux and Windows. Table S5 gives the complete command lines.

### Timing

We used the production input for replica 1 of the GLP-1-bound wild-type receptor from the study by Kato et al. [24]. The membrane system contained 31,884 OPC waters, 240 palmitoyl-oleoyl phosphatidylcholine lipids and 421 CMAP interactions. Fourteen configurations ran in three randomized blocks on an RTX 3080 and an RTX 2080 Ti (84 runs), both installed in one workstation with an Intel Core i7-13700K processor. We timed the final 75,000 steps of each 100,000-step (0.2 ns) run. Elapsed times were recorded in whole seconds, giving a resolution of 0.013 ms per step. For each device, we divided the mean of the three within-block savings at each change by the mean step time of the initial Optimized configuration with CPU update and CPU CMAP. The share attributed to GPU update uses the sum of the three positive changes in ms per step. Table S3 gives the per-device step-time ratios and the throughput of further configurations.

## Acknowledgments

We thank the GROMACS developers for the open-source software on which this work builds, and Lukas Müllender in particular for making the GPU virtual-site prototypes publicly available. We also thank the developers of the ff19SB port to GROMACS and of the OPC water model.

## Author contributions

Takashi Kato designed the study, performed the simulations and analyses, and wrote the manuscript. Nobuhiro Shojima and Toshimasa Yamauchi advised on the study and helped verify the results.

## Data and code availability

The Supplementary Methods give the full simulation settings, the fold-detection procedure and the benchmark configurations, with Figures S1 and S2 and Tables S1–S5. Supplementary Data 1 contains the numerical data for all figures and tables. The modified GROMACS source, its unit tests and the GLP-1R benchmark settings are available at https://github.com/toropippi/gromacs-opcfast/tree/opcfast-paper-v1. Complete trajectories are available from the corresponding author on request. The GROMACS patches retain the GNU Lesser General Public License, version 2.1 or later.

## Funding and competing interests

The work received no external funding. The authors declare no competing interests.

## AI assistance

AI tools assisted with literature searches, review of GROMACS source code and public development records, analysis code and writing. The authors are responsible for the scientific verification and final text.

## Notes

https://github.com/toropippi/gromacs-opcfast/tree/opcfast-paper-v1

### Competing Interest Statement

The authors have declared no competing interest.

## References

[1] Karplus M, McCammon JA. Molecular dynamics simulations of biomolecules. Nat Struct Biol. 2002;9:646–652. doi:10.1038/nsb0902-646.

[2] Dror RO, Dirks RM, Grossman JP, Xu H, Shaw DE. Biomolecular Simulation: A Computational Microscope for Molecular Biology. Annu Rev Biophys. 2012;41:429–452. doi:10.1146/annurev-biophys-042910-155245.

[3] Hollingsworth SA, Dror RO. Molecular Dynamics Simulation for All. Neuron. 2018;99:1129–1143. doi:10.1016/j.neuron.2018.08.011.

[4] Shaw DE, Maragakis P, Lindorff-Larsen K, et al. Atomic-Level Characterization of the Structural Dynamics of Proteins. Science. 2010;330:341–346. doi:10.1126/science.1187409.

[5] Lindorff-Larsen K, Piana S, Dror RO, Shaw DE. How Fast-Folding Proteins Fold. Science. 2011;334:517–520. doi:10.1126/science.1208351.

[6] Latorraca NR, Venkatakrishnan AJ, Dror RO. GPCR Dynamics: Structures in Motion. Chem Rev. 2017;117:139–155. doi:10.1021/acs.chemrev.6b00177.

[7] De Vivo M, Masetti M, Bottegoni G, Cavalli A. Role of Molecular Dynamics and Related Methods in Drug Discovery. J Med Chem. 2016;59:4035–4061. doi:10.1021/acs.jmedchem.5b01684.

[8] Friedrichs MS, Eastman P, Vaidyanathan V, et al. Accelerating molecular dynamic simulation on graphics processing units. J Comput Chem. 2009;30:864–872. doi:10.1002/jcc.21209.

[9] Salomon-Ferrer R, Götz AW, Poole D, Le Grand S, Walker RC. Routine Microsecond Molecular Dynamics Simulations with AMBER on GPUs. 2. Explicit Solvent Particle Mesh Ewald. J Chem Theory Comput. 2013;9:3878–3888. doi:10.1021/ct400314y.

[10] Eastman P, Swails J, Chodera JD, et al. OpenMM 7: Rapid development of high performance algorithms for molecular dynamics. PLoS Comput Biol. 2017;13:e1005659. doi:10.1371/journal.pcbi.1005659.

[11] Páll S, Zhmurov A, Bauer P, Abraham M, Lundborg M, Gray A, Hess B, Lindahl E. Heterogeneous parallelization and acceleration of molecular dynamics simulations in GROMACS. J Chem Phys. 2020;153:134110. doi:10.1063/5.0018516.

[12] Onufriev AV, Izadi S. Water models for biomolecular simulations. WIREs Comput Mol Sci. 2018;8:e1347. doi:10.1002/wcms.1347.

[13] Jorgensen WL, Chandrasekhar J, Madura JD, Impey RW, Klein ML. Comparison of simple potential functions for simulating liquid water. J Chem Phys. 1983;79:926–935. doi:10.1063/1.445869.

[14] Horn HW, Swope WC, Pitera JW, Madura JD, Dick TJ, Hura GL, Head-Gordon T. Development of an improved four-site water model for biomolecular simulations: TIP4P-Ew. J Chem Phys. 2004;120:9665–9678. doi:10.1063/1.1683075.

[15] Abascal JLF, Vega C. A general purpose model for the condensed phases of water: TIP4P/2005. J Chem Phys. 2005;123:234505. doi:10.1063/1.2121687.

[16] Izadi S, Anandakrishnan R, Onufriev AV. Building Water Models: A Different Approach. J Phys Chem Lett. 2014;5:3863–3871. doi:10.1021/jz501780a.

[17] Shabane PS, Izadi S, Onufriev AV. General Purpose Water Model Can Improve Atomistic Simulations of Intrinsically Disordered Proteins. J Chem Theory Comput. 2019;15:2620–2634. doi:10.1021/acs.jctc.8b01123.

[18] Tian C, et al. ff19SB: Amino-Acid-Specific Protein Backbone Parameters Trained against Quantum Mechanics Energy Surfaces in Solution. J Chem Theory Comput. 2020;16:528–552. doi:10.1021/acs.jctc.9b00591.

[19] Abraham MJ, Murtola T, Schulz R, Páll S, Smith JC, Hess B, Lindahl E. GROMACS: High performance molecular simulations through multi-level parallelism from laptops to supercomputers. SoftwareX . 2015;1–2:19–25. doi:10.1016/j.softx.2015.06.001.

[20] Feenstra KA, Hess B, Berendsen HJC. Improving efficiency of large time-scale molecular dynamics simulations of hydrogen-rich systems. J Comput Chem. 1999;20:786–798.

[21] GROMACS developers. Getting good performance from mdrun. GROMACS 2026.3 documentation. https://manual.gromacs.org/2026.3/user-guide/mdrun-performance.html.

[22] Drucker DJ.Mechanisms of Action and Therapeutic Application of Glucagon-like Peptide-1. Cell Metab. 2018;27:740–756. doi:10.1016/j.cmet.2018.03.001.

[23] Zhang Y, Sun B, Feng D, et al. Cryo-EM structure of the activated GLP-1 receptor in complex with a G protein. Nature. 2017;546:248–253. doi:10.1038/nature22394.

[24] Kato T, Inomata N, Serizawa K, et al. GLP-1R R131Q links long-range conformational remodeling to cellular stress phenotypes. J Mol Graph Model. 2026;146:109445. doi:10.1016/j.jmgm.2026.109445.

[25] Müllender L. Implement virtual sites on GPU. GROMACS merge request 5977, opened May 5, 2026. https://gitlab.com/gromacs/gromacs/-/merge_requests/5977 (accessed November 19, 2026).

[26] Müllender L. Enable GPU update for type 1, 2 and 3 VSites. GROMACS merge request 6119, opened July 10, 2026, merged November 17, 2026. https://gitlab.com/gromacs/gromacs/-/merge_requests/6119 (accessed November 20, 2026).

[27] Goldberg D. What every computer scientist should know about floating-point arithmetic. ACM Comput Surv. 1991;23:5–48. doi:10.1145/103162.103163.

[28] Le Grand S, Götz AW, Walker RC. SPFP: Speed without compromise—A mixed precision model for GPU accelerated molecular dynamics simulations. Comput Phys Commun. 2013;184:374–380. doi:10.1016/j.cpc.2012.09.022.

[29] Braxenthaler M, Unger R, Auerbach D, Given JA, Moult J. Chaos in protein dynamics. Proteins. 1997;29:417–425. doi:10.1002/(SICI)1097-0134(199712)29:4<417::AID-PROT2>3.0.CO;2-5.

[30] Simmerling C, Strockbine B, Roitberg AE. All-Atom Structure Prediction and Folding Simulations of a Stable Protein. J Am Chem Soc. 2002;124:11258–11259. doi:10.1021/ja0273851.

[31] Neidigh JW, Fesinmeyer RM, Andersen NH. Designing a 20-residue protein. Nat Struct Biol. 2002;9:425–430. doi:10.1038/nsb798.

[32] Qiu L, Pabit SA, Roitberg AE, Hagen SJ. Smaller and Faster: The 20-Residue Trp-Cage Protein Folds in 4 µs. J Am Chem Soc. 2002;124:12952–12953. doi:10.1021/ja0279141.

[33] Paschek D, Day R, García AE. Influence of water–protein hydrogen bonding on the stability of Trp-cage miniprotein. A comparison between the TIP3P and TIP4P-Ew water models. Phys Chem Chem Phys. 2011;13:19840–19847. doi:10.1039/C1CP22110H.

[34] Singh B, Martínez-Noa Y, Perez A. How well do molecular dynamics force fields model peptides: a systematic benchmark across diverse folding behaviors. J Phys Chem B. 2026;130:4344–4357. doi:10.1021/acs.jpcb.6c01176.

[35] Miletić V, Wieczór M, Simmerling C, Rampp M, de Groot BL, Kutzner C, Gapsys V. Force Fields and Simulation Engines: Lessons Learned from Porting Amber ff14SB and ff19SB to GROMACS. J Chem Theory Comput. 2026;22:11142–11152. doi:10.1021/acs.jctc.6c01592.

[36] Kabsch W, Sander C. Dictionary of protein secondary structure: Pattern recognition of hydrogen-bonded and geometrical features. Biopolymers. 1983;22:2577–2637. doi:10.1002/bip.360221211.

