## Supplementary Information for "GPU handling of OPC water in GROMACS: validation by protein folding and a 1.6-fold speed-up"

This file contains the Supplementary Methods, Figures S1 and S2, Tables S1–S5 and the Supplementary References.

### Supplementary Methods

#### Trp-cage simulations and water measurements (Figure 1, Figure S1 and Table S1)

We simulated the 20-residue Trp-cage miniprotein (TC5b) with the ff19SB protein force field [1] and four-site Optimal Point Charge (OPC) water [2], using the GROMACS port of ff19SB [3]. GROMACS 2026.3 (Reference) and the modified build `opcfast20260809g` (Optimized) read identical compiled inputs within each matched pair. Each system contained the peptide with charged termini, one neutralizing chloride and 9,300 waters. The starting peptide was extended in a rhombic dodecahedral box with a nominal 1.2 nm solute-to-wall separation.

After energy minimization, five replicas received independent initial velocities. Each underwent 0.5 ns of position-restrained equilibration at constant volume, 1 ns with restraints at constant pressure, then 20 ns at constant pressure with restraints released. Each equilibrated structure and its velocities started one 5 $\mu$s trajectory per build. Production used a 2 fs leapfrog step, velocity-rescaling temperature coupling [4] at 300 K ($\tau_T = 1$ ps) and isotropic Parrinello–Rahman pressure coupling [5] at 1 bar ($\tau_P = 5$ ps). Protein bonds were constrained with LINCS [6] (order four, one iteration) and water geometry with SETTLE [7]. The real-space cutoff for particle-mesh Ewald electrostatics [8] and the Lennard–Jones cutoff were 1.0 nm. We used fourth-order mesh interpolation, analytical dispersion corrections and a 150-step neighbor-list interval.

**Charge-site positions and forces.** The OPC negative charge lies at a massless site M. We reconstructed M from the final oxygen and hydrogen coordinates for every water in each replica, using the nearest periodic O–H displacement vectors $\mathbf{d}_1$ and $\mathbf{d}_2$ and the coefficient $a$ from the simulation input:

$$\mathbf{r}_M = \mathbf{r}_O + a(\mathbf{d}_1 + \mathbf{d}_2), \qquad a \simeq 0.1477223.$$

For each endpoint we report the root-mean-square distance between reconstructed and stored sites, and we compared the five values of each build with Welch's $t$-test. The force on M is distributed to O, H1 and H2 with weights $1 - 2a$, $a$ and $a$, respectively. We compared unrounded Cartesian forces from separate runs on a common initial configuration. For each atom selection we report the root-mean-square length of the force-difference vectors (Optimized minus Reference) and of the Reference force vectors. The relative force difference is the Euclidean norm of the difference divided by the norm of the Reference forces. The per-molecule comparison uses the sum over its four sites.

**Solvent structure.** At each endpoint we binned intermolecular O–O and H–M distances up to 0.55 nm in 0.005 nm bins, including periodic images. Counts were divided by the number of centers, the shell volume and the number density of the other 9,299 waters. Each replica contributed equally to its group mean. For lysine 8 we counted water oxygens within 0.35 nm of its side-chain nitrogen and averaged the cosine of the angle between the nitrogen-to-oxygen and oxygen-to-M vectors. Densities are time averages from the full trajectories.

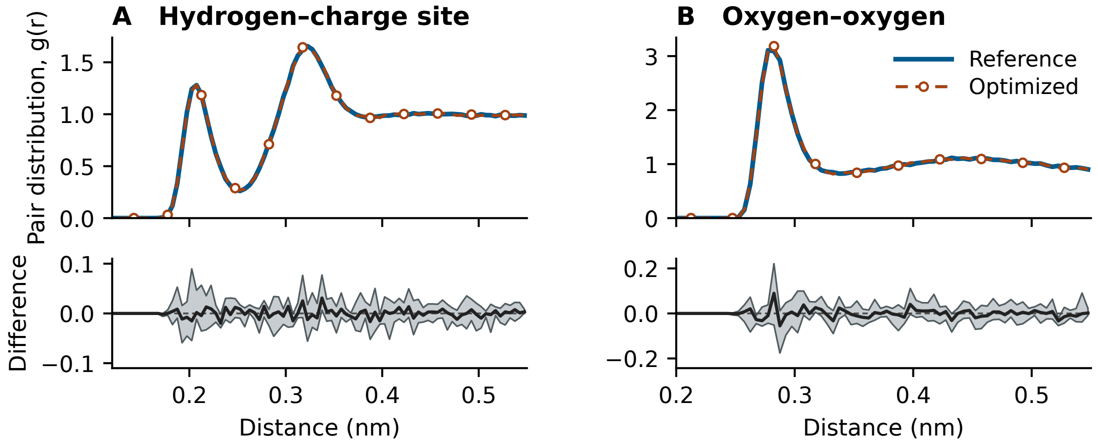


Figure S1: Water pair distributions at 5 $\mu$s. Upper panels show five-endpoint means for Reference (solid) and Optimized (dashed with circles). Lower panels show matched differences (Optimized minus Reference), their mean (solid), range (shaded) and zero (dashed).

Table S1: Solvent structure, hydration and density, averaged over five replicas. The Lys8 shell includes water oxygens within 0.35 nm of its side-chain nitrogen. Positive cosines point the O–M vector away from the nitrogen.

| Quantity | Reference | Optimized | Sampling |
|---|---|---|---|
| Other M sites per H within 0.24 nm | 0.8445 | 0.8446 | Endpoints |
| Other O atoms per O within 0.35 nm | 5.167 | 5.183 | Endpoints |
| Lys8 shell waters | 4.0 | 5.2 | Endpoints |
| Lys8 shell orientation cosine | 0.62 | 0.49 | Endpoints |
| Solution density (kg $m^{-3}$) | 1000.48 | 1000.48 | 0–5 $\mu$s |

The first intermolecular H–M peak fell at 0.205–0.210 nm in both builds, and the oxygen–oxygen peaks occupied adjacent bins (Figure S1). The water–water quantities in Table S1 agree between the builds. The Lys8 values are single-frame measurements around one side chain and depend on the conformation of the peptide, which differed between the endpoints: two Reference and four Optimized endpoints lay within 0.25 nm of the native structure by C$\alpha$ root-mean-square deviation. They are therefore not a comparison of the builds.

**Numerical controls.** In separate 50 ps constant-energy Trp-cage tests, the magnitude of the fitted energy drift increased as the time step decreased from 2 to 0.5 fs in both builds. Those runs did not establish time-step convergence.

#### Trp-cage folding analysis (Figures 2–3, Table 1, Figure S2 and Table S2)

We reconstructed whole peptide molecules across periodic boundaries and fitted the C$\alpha$ atoms to model 1 of the 1L2Y solution NMR structure with equal weights. We screened all 50,001 frames per trajectory at 100 ps intervals. A frame was classified as native-like when it satisfied all four criteria:

1. C$\alpha$ root-mean-square deviation (RMSD) from the reference structure $\leq 0.25$ nm.
2. At least 75% of the 19 native C$\alpha$ contacts present. Native pairs are separated by at least three residues in sequence and by less than 0.65 nm in the reference structure. A pair was counted as present when its distance in a frame was $\leq 0.65$ nm.
3. Minimum heavy-atom distances from the Trp6 indole to the side chains of Pro12 and Pro18 both $\leq 0.42$ nm. The proline selection comprises CB, CG and CD.
4. At least five of residues 2–8 assigned to an $\alpha$-helix by the GROMACS implementation of the Kabsch–Sander secondary-structure algorithm [9].

We identified candidate persistent events when at least 80% of frames in a 10 ns window were native-like. Overlapping candidates were merged, extended by about 5 ns on each side and examined in time order at 10 ps intervals, up to 120 ns per candidate. Confirmation required a frame with RMSD $\leq 0.15$ nm after fitting the C$\alpha$ atoms of residues 3–19 and the heavy atoms of Trp6 from CB through the indole ring. Entry time is the first frame satisfying that criterion in the confirmed interval. For the separate 100 ns persistence measure we selected a high-occupancy window on the coarse grid and checked it at 10 ps intervals. For Figure 3 we took, from every trajectory with a confirmed persistent event, the last frame classified as native-like in the 100 ps screening, made the peptide whole across periodic boundaries and fitted its C$\alpha$ atoms to model 1 of 1L2Y. The structures were drawn with open-source PyMOL. For Figure 3B and Table 1 we used every frame classified as native-like in the trajectories with a confirmed persistent event (three Reference and five Optimized trajectories; 43,482 and 88,114 frames). Distributions pool the frames of each build. Their overlap is the sum over bins of the smaller of the two fractions. Table 2 averages each quantity over the native-like frames of one trajectory and reports the mean and sample standard deviation of these trajectory means. Because native-like frames are defined by thresholds on the same quantities, the distributions are bounded by those thresholds. Numbers of folded trajectories were compared with Fisher's exact test, first-entry times with the log-rank test, treating trajectories without a persistent event as censored at 5 $\mu$s, and full-trajectory occupancies with Welch's $t$-test.

Table S2: Trp-cage folding by starting input. Full occupancy uses every 100 ps frame across 5 $\mu$s. Entry and selected-window occupancy use the finer 10 ps sampling described above. A dash denotes no confirmed event or window.

| Build | Replica | Full occupancy (%) | Entry (ns) | Selected 100 ns (%) |
|---|---|---|---|---|
| Reference | r1 | 29.33 | 251.32 | 88.34 |
| Reference | r2 | 0.88 | — | — |
| Reference | r3 | 34.62 | 846.90 | 88.16 |
| Reference | r4 | 0.00 | — | — |
| Reference | r5 | 23.01 | 3416.33 | 88.04 |
| Optimized | r1 | 51.43 | 1408.02 | 88.11 |
| Optimized | r2 | 29.04 | 2616.23 | 87.41 |
| Optimized | r3 | 6.44 | 3261.29 | 86.28 |
| Optimized | r4 | 54.07 | 316.88 | 88.77 |
| Optimized | r5 | 35.24 | 2772.35 | 88.61 |

Full-trajectory occupancies of the two builds were compared with Welch's $t$-test, treating the five runs of each build as independent samples. Runs that share a starting structure diverge within picoseconds, so the pairing carries no information about when a run folds.

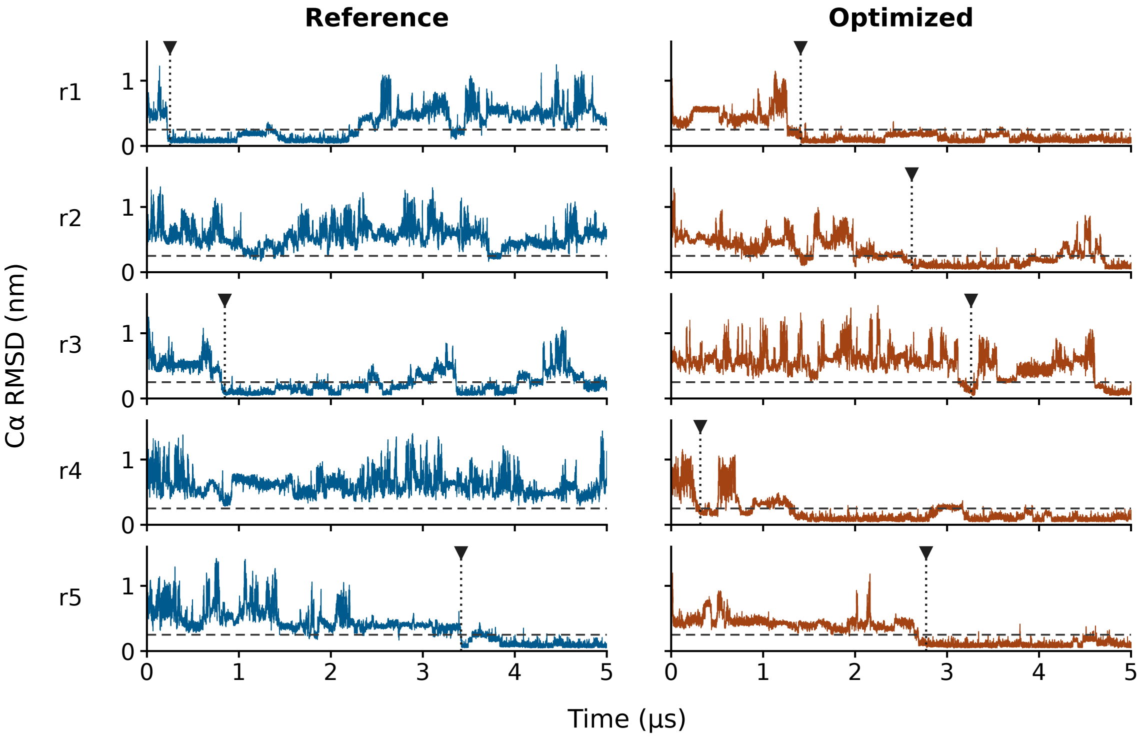


Figure S2: Trp-cage C$\alpha$ root-mean-square deviation (RMSD) from NMR model 1 of 1L2Y over 5 $\mu$s, sampled every 100 ps. Each row shows two runs from the same starting input. Horizontal dashed lines mark the 0.25 nm threshold. Triangles and vertical dotted lines mark the first confirmed native-like event.

#### Bulk water and receptor hydration (Figure 4 and Table 2)

**Pure water.** Boxes of 9,300 TIP3P or OPC water molecules, with the parameters of the ff19SB port, were energy-minimized, equilibrated for 0.5 ns and simulated for 10 ns with the settings in Table S4. OPC Reference and OPC Optimized read the same compiled production input, so they started from identical coordinates and velocities. TIP3P ran on the RTX 2080 Ti and OPC Optimized on the RTX 3080 of the benchmark workstation. OPC Reference ran on a second workstation with an AMD Ryzen 9 9950X processor and an RTX 5090, using GROMACS 2026.3 built from the official release with CUDA 12.9.

Each 10-ns trajectory was cut into five 2-ns blocks. For each block we computed four properties with the unmodified GROMACS 2026.3 analysis tools:

- density, as the block mean (`gmx energy`);
- the self-diffusion coefficient, from a linear fit to the mean-square displacement of the oxygens between lag times of 50 and 500 ps, with time origins every 20 ps (`gmx msd`);
- the static dielectric constant, from the fluctuation of the total dipole moment at 298.15 K (`gmx dipoles`);
- $g_{OO}$, in 0.002 nm bins from frames 10 ps apart (`gmx rdf`); the heights of the first and second peaks are its maxima before and after the first minimum.

**GLP-1R hydration.** The starting coordinates and velocities of the published simulation of replica 1 of the GLP-1-bound wild-type receptor were written with nine decimals and recompiled with the published settings, changed only so that all atoms were saved every 10 ps. Both builds read this one input and ran for 8 ns on the RTX 3080 with the options of Table S5 (tuned Reference; Optimized with all features). We analysed the 800 frames after the common starting frame with the following definitions:

- *First hydration shell of a residue*: water molecules whose oxygen lies within 0.35 nm of any nitrogen or oxygen atom of the residue, under periodic boundary conditions. Residue means are averages over all frames.
- *Bridging water*: a water in the first shell of at least one of GLP-1 residues His7–Asp15 and of at least one receptor residue.
- *Residence curve*: $C(t) = \langle h(0)h(t)\rangle / \langle h\rangle$, where $h$ is 1 when a water is in the first shell of His7–Asp15 and 0 otherwise.

Block statistics used four consecutive 2-ns blocks per build.

#### GLP-1 receptor membrane benchmark (Figure 5, Table 3 and Table S3)

The performance input was the production input of replica 1 of the GLP-1-bound wild-type receptor from our previous study [10]. It contained glucagon-like peptide-1 (GLP-1) bound to its receptor in a bilayer, with 166,826 atoms, 240 phosphatidylcholine lipids and 31,884 OPC waters. The protein had 421 backbone dihedral correction-map interactions (CMAP). We ran 14 configurations in three randomized blocks on each of two graphics processing units (GPUs), an RTX 3080 and an RTX 2080 Ti. Both GPUs were installed in one Windows workstation with an Intel Core i7-13700K processor (16 cores, 24 threads), and each run used one GPU. Both builds were compiled with MSVC 19.44 and

CUDA 12.6. Each run advanced 100,000 steps at 2 fs. Timing began after 25,000 steps. The 84 timed runs are tabulated individually in Supplementary Data 1.

**Run options.** Table S5 lists the command lines. Both builds offloaded nonbonded interactions and particle-mesh Ewald (PME) electrostatics to the GPU. The Optimized build detects OPC water in the input and needs no option of its own: with `-bonded gpu -update gpu`, OPC site construction, force redistribution and CMAP run on the GPU, and CUDA Graphs are used unless the environment variable `GMX_DISABLE_CUDA_GRAPH` is set. The comparison with tuned Reference used automatic PME tuning for both builds; PME settings were held fixed (`-notunepme`) when timing successive changes.

The options and the variable do not depend on the operating system. The variable is set with `export` in a Linux shell and with `set` in the Windows command prompt. The source builds with the standard GROMACS CMake procedure on both systems, and the configuration command is given in the repository. Besides the Windows builds used for the measurements reported here, build `opcfast20260809g` has been compiled on Ubuntu 20.04 with GCC 11.5 and CUDA 12.9, where it runs production simulations with the same options.

**Incremental comparisons.** Starting from the Optimized build with CPU coordinate updates and CPU CMAP, we enabled each change in Table S3 in sequence. For each block we subtracted the time per step after a change from the time before it. Figure 5 expresses the mean saving as a percentage of the initial CPU-update time on that GPU (1.933 and 2.058 ms per step on the RTX 3080 and RTX 2080 Ti). Bars average the two device percentages. For the share attributable to GPU update, the denominator is the total time saved by the first three changes on each device.

Table S3: GLP-1R performance. Step-time ratios divide the mean before each change in Figure 5 by the mean after it. Throughput is the mean ± sample standard deviation of three timed runs, in ns/day.

| Change (step-time ratio) | RTX 3080 | RTX 2080 Ti |
|---|---|---|
| GPU coordinate updates, constraints and OPC sites | 1.667 | 1.684 |
| GPU backbone correction (CMAP) | 1.083 | 1.007 |
| Fuse water-force redistribution with force reduction | 1.021 | 1.011 |
| Enable CUDA Graphs | 0.944 | 0.964 |
| **Configuration (ns/day)** | **RTX 3080** | **RTX 2080 Ti** |
| Tuned Reference (PME tuning on) | $94.6 \pm 0.0$ | $88.0 \pm 0.9$ |
| Optimized, graphs off (fixed PME) | $164.8 \pm 1.2$ | $144.0 \pm 0.0$ |
| Optimized, graphs on (fixed PME) | $155.5 \pm 1.1$ | $138.9 \pm 0.9$ |
| Optimized, graphs on (PME tuning on) | $156.8 \pm 1.1$ | $138.4 \pm 1.7$ |

**Coordinate transfers.** We profiled three configurations twice per device with Nsight Systems: CPU update and CPU CMAP, GPU update and CPU CMAP, and GPU update and GPU CMAP. Fusion and CUDA Graphs were disabled for these traces. Each trace covered 8,000 steps. We measured host-to-device traffic across 5,500 intervals between successive electrostatic mesh-gather operations. Moving the update to the GPU reduced the traffic from about 4.11 to 2.19 MB per interval on both devices. A full coordinate vector occupies 2.00 MB, consistent with removing one coordinate upload per regular step.

Table S4: Settings of the pure-water simulations (Figure 4A and Table 2 of the main text). All stages used the same settings unless a stage is named.

| Setting | Value |
|---|---|
| System | 9,300 water molecules in a cubic box of initially 6.6 nm |
| Water geometry | rigid |
| Time step | 2 fs |
| Thermostat | velocity rescaling, 298.15 K, time constant 1 ps |
| Barostat, equilibration | stochastic cell rescaling, 1 bar, time constant 2 ps |
| Barostat, production | Parrinello–Rahman, 1 bar, time constant 5 ps |
| Compressibility | $4.5 \times 10^{-5}$ bar$^{-1}$ |
| Electrostatics | particle-mesh Ewald, grid spacing 0.12 nm, fourth-order interpolation |
| Cut-offs | 1.0 nm for Coulomb and Lennard-Jones interactions, potential shift |
| Dispersion correction | energy and pressure |
| Equilibration | 0.5 ns from velocities drawn at 298.15 K |
| Production | 10 ns, coordinates and energies saved every 1 ps |

Table S5: Command lines of the GLP-1R runs. All configurations shared `gmx mdrun -s md_0_1.tpr -deffnm md -nsteps 100000 -resetstep 25000 -noconfout -gpu_id 0 -ntmpi 1 -ntomp 6 -nb gpu -pme gpu -pmefft gpu -nstlist 150`; the RTX 2080 Ti runs used `-ntomp 24`. The columns give the options and the environment variable that differ.

| Configuration | Binary | Additional options | Environment variable |
|---|---|---|---|
| Tuned Reference | GROMACS 2026.3 | `-bonded cpu -update cpu -tunepme` | none |
| Optimized, all features | `opcfast` build | `-bonded gpu -update gpu -tunepme` | none |
| Optimized, fastest | `opcfast` build | `-bonded gpu -update gpu -notunepme` | `GMX_DISABLE_CUDA_GRAPH=1` |

### Supplementary References


[1] Tian C, et al. ff19SB: Amino-Acid-Specific Protein Backbone Parameters Trained against Quantum Mechanics Energy Surfaces in Solution. *J Chem Theory Comput*. 2020;16:528–552. doi:10.1021/acs.jctc.9b00591.

[2] Izadi S, Anandakrishnan R, Onufriev AV. Building Water Models: A Different Approach. *J Phys Chem Lett*. 2014;5:3863–3871. doi:10.1021/jz501780a.

[3] Miletić V, Wieczór M, Simmerling C, Rampp M, de Groot BL, Kutzner C, Gapsys V. Force Fields and Simulation Engines: Lessons Learned from Porting Amber ff14SB and ff19SB to GROMACS. *J Chem Theory Comput*. 2026;22:11142–11152. doi:10.1021/acs.jctc.6c01592.

[4] Bussi G, Donadio D, Parrinello M. Canonical sampling through velocity rescaling. *J Chem Phys*. 2007;126:014101. doi:10.1063/1.2408420.

[5] Parrinello M, Rahman A. Polymorphic transitions in single crystals: A new molecular dynamics method. *J Appl Phys*. 1981;52:7182–7190. doi:10.1063/1.328693.

[6] Hess B, Bekker H, Berendsen HJC, Fraaije JGEM. LINCS: A linear constraint solver for molecular simulations. *J Comput Chem*. 1997;18:1463–1472.

[7] Miyamoto S, Kollman PA. Settle: An analytical version of the SHAKE and RATTLE algorithm for rigid water models. *J Comput Chem*. 1992;13:952–962. doi:10.1002/jcc.540130805.

[8] Essmann U, Perera L, Berkowitz ML, Darden T, Lee H, Pedersen LG. A smooth particle mesh Ewald method. *J Chem Phys*. 1995;103:8577–8593. doi:10.1063/1.470117.

[9] Kabsch W, Sander C. Dictionary of protein secondary structure: Pattern recognition of hydrogen-bonded and geometrical features. *Biopolymers*. 1983;22:2577–2637. doi:10.1002/bip.360221211.

[10] Kato T, Inomata N, Serizawa K, et al. GLP-1R R131Q links long-range conformational remodeling to cellular stress phenotypes. *J Mol Graph Model*. 2026;146:109445. doi:10.1016/j.jmgm.2026.109445.
